# miR-106b-responsive gene landscape identifies regulation of Kruppel-like factor family

**DOI:** 10.1101/088229

**Authors:** Cody J. Wehrkamp, Sathish Kumar Natarajan, Ashley M. Mohr, Mary Anne Phillippi, Justin L. Mott

**Author notes:** Address for Correspondence: Justin L. Mott, MD, PhD Associate Professor Department of Biochemistry and Molecular Biology 985870 Nebraska Medical Center Omaha, NE 68198-5870.

## Abstract

MicroRNA dysregulation is a common feature of cancer and due to the promiscuity of microRNA binding this can result in a wide array of genes whose expression is altered. miR-106b is an oncomiR overexpressed in cholangiocarcinoma and its upregulation in this and other cancers often leads to repression of anti-tumorigenic targets. The goal of this study was to identify the miR-106b-regulated gene landscape in cholangiocarcinoma cells using a genome-wide, unbiased mRNA analysis. Through RNA-Seq we found 112 mRNAs significantly repressed by miR-106b. The majority of these genes contain the specific miR-106b seed-binding site. We have validated 11 genes from this set at the mRNA level and demonstrated regulation by miR-106b of five proteins. Combined analysis of our miR-106b-regulated mRNA data set plus published reports indicate that miR-106b binding is anchored by G:C pairing in and near the seed. Novel targets Kruppel-like factor 2 (KLF2) and KLF6 were verified both at the mRNA and at the protein level. Further investigation showed regulation of four other KLF family members by miR-106b. We have discovered coordinated repression of several members of the KLF family by miR-106b that may play a role in cholangiocarcinoma tumor biology.

**List of abbreviations**
MTT: 3-(4,5-dimethylthiazol-2-yl)-2,5-diphenyltetrazolium bromide
DAPI: 4’,6-diamidine-2’-phenylindole dihydrochloride
CLASH: Crosslinking, ligation, and sequencing of hybrids
DR5: Death receptor-5
DMEM: Dulbecco’s Modified Eagle’s Medium
FDR: False discovery rate
FBS: Fetal bovine serum
KLF: Kruppel-like factor
LNA: Locked-nucleic acid
qRT-PCR: Quantitative reverse-transcription PCR
SDS-PAGE: Sodium dodecylsulfate-polyacrylamide gel electrophoresis
UTR: Untranslated region

## Introduction

miR-106b has been established as an oncogenic microRNA with increased expression in many cancers including cholangiocarcinoma [12345], prostate [6], gastric [7], and hepatocellular carcinoma [8]. Some functions of miR-106b include increased proliferation by miR-106b-mediated reduction of the transcription factor E2F1 [9] and the tumor suppressor RB1 [10]. Additionally, miR-106b increased migration through targeting of the phosphatase PTEN [11], and reduced apoptosis by regulating the BH3-containing protein Bim [12].

MicroRNAs function to dampen the expression of their targets [13,14,15]. This is accomplished both through reduction in mRNA transcript level as well as by repression of translation. While both mechanisms contribute to reduced target protein expression, the effect of mRNA reduction may dominate [16], in part because each message is used as a transcript for synthesis of many protein molecules; still, the magnitude of change in mRNA expression is low. A focused description of the binding characteristics and targets of individual microRNAs is feasible and desirable.

The seed region of a microRNA is a 7-8 nucleotide sequence at the 5’ end of the microRNA which is vital for complementary binding to its mRNA target. The complementary seed-binding site is more or less conserved among targets, allowing for broad transcript-expression dampening effects from a single microRNA. Seed-binding sites are highly enriched in the 3’ untranslated region (UTR) of transcripts [17]. There is also evidence that supports non-canonical, non-seed mRNA target interactions between microRNAs and their targets [18, 19]. The traditional seed for miR-106b is located at nucleotides 1-8 on its 5’ end, corresponding to the microRNA sequence 5’UAAAGUGC.

In this study, we sought to define the genome-wide target set of miR-106b and found that miR-106b binding relied on sequences between bases 2-10, tolerating a G:U wobble in the seed region. We found 112 transcripts that were regulated at the mRNA level in cholangiocarcinoma cells. Finally, we demonstrated that members of the KLF family of transcription factors are coordinately repressed by miR-106b. Comparison of our data in cholangiocarcinoma cells with data in Flp-In T-REx 293 human embryonic kidney-derived cells [19] revealed that the microRNA-responsive gene set is largely cell-type specific.

## Materials and Methods

### Cell lines

Human cholangiocarcinoma cell lines were previously derived from a female patient with metastatic gallbladder cancer, Mz-ChA-1 [20], or a male patient with combined histologic features of hepatocellular carcinoma and cholangiocarcinoma, KMCH [21]. BDEneu rat cholangiocarcinoma cells were derived from primary Fisher 344 rat cholangiocytes [22]. H69 cells are a non-malignant immortalized cholangiocyte cell line. Cholangiocarcinoma cells were grown in Dulbecco’s Modified Eagle’s Medium (DMEM) supplemented with 10% fetal bovine serum (FBS), insulin (0.5 μg/mL) and G418 (50 μg/mL). H69 cells were grown in DMEM supplemented with 10% FBS, insulin (5 μg/mL), adenine (24.3 μg/mL), epinephrine (1 ug/mL), tri-iodothyronine (T3)-transferrin (T), [T3-2.23 ng/ml, T-8.19 μg/mL], epidermal growth factor (9.9 ng/mL) and hydrocortisone (5.34 μg/mL).

Cells were transfected either with mature miR-106b mimic (Life Technologies), locked-nucleic acid (LNA) antagonist to miR-106b, LNA-106b (Exiqon), or LNA Negative Control A (Exiqon) for 24-48 hours using Lipofectamine RNAiMAX at a final oligonucleotide concentration of 20 nM. miR-106b levels were quantified by TaqMan Small RNA assay (Life Technologies).

### RNA-Seq

High quality total RNA was extracted from Mz-ChA-1 cells transfected with either miR-106b (n=3) or LNA-106b (n=3) for RNA-Seq using RNeasy Mini Kit (Qiagen). The quality of the RNA was confirmed by RNA Integrity Analysis using an Agilent Bioanalyzer. RNA sequencing libraries were prepared by the UNMC Next Generation Sequencing Core Facility beginning with 1 μg of total RNA and a TruSeq RNA Prep V2 (Illumina Inc, San Diego, CA). Samples were sequenced using the Illumina HiSeq 2000 system with 100 bp paired-end reads. An average of 76.99 million reads per sample were collected (range 58.99-87.00 million).

Sequences were analyzed by the Bioinformatics and Systems Biology Core at the University of Nebraska Medical Center (UNMC) using the Tuxedo protocol [23]. Reads were mapped to the human genome using Top Hat. Transcripts were assigned by Cuff links and the transcriptome defined using Cuffmerge. Finally, differential expression between miR-106b and LNA-106b was calculated with Cuffdiff.

Using a false discovery rate (FDR)-corrected p value, genome-wide transcripts were ranked from most significant to least. This ranked list of genes was submitted to the SylArray online server to detect enrichment of microRNA seed-binding sites [24].

### miR-106b binding site analysis

RefSeq sequences for the 129 significant genes (112 decreased and 17 increased) were collected in Fasta format and analyzed by direct search for the known miR-106b binding site as well as subjected to k-mer analysis to generate a k table of all possible 7-mer and 8-mer sequences and their frequency in this gene set. Several genes had more than one RefSeq sequence, so the final size of this Fasta file was 191 transcripts. For comparison, 1,000 random sets of 191 transcripts each from the RefSeq database were generated.

RNAHybrid [25] was used to identify the single best miR-106b predicted binding site for each decreased transcript based on minimum free energy hybridization. The resulting miR-106b-binding sites were manually curated for discovery of ungapped motifs using MEME Suite [26].

### Quantitative reverse-transcription PCR

Total RNA for qRT-PCR was isolated using TRIzol Reagent (Life Technologies) and quantified using SYBR Green DNA binding (Roche). Primer pairs are listed in Table 1.

**Table 1:**
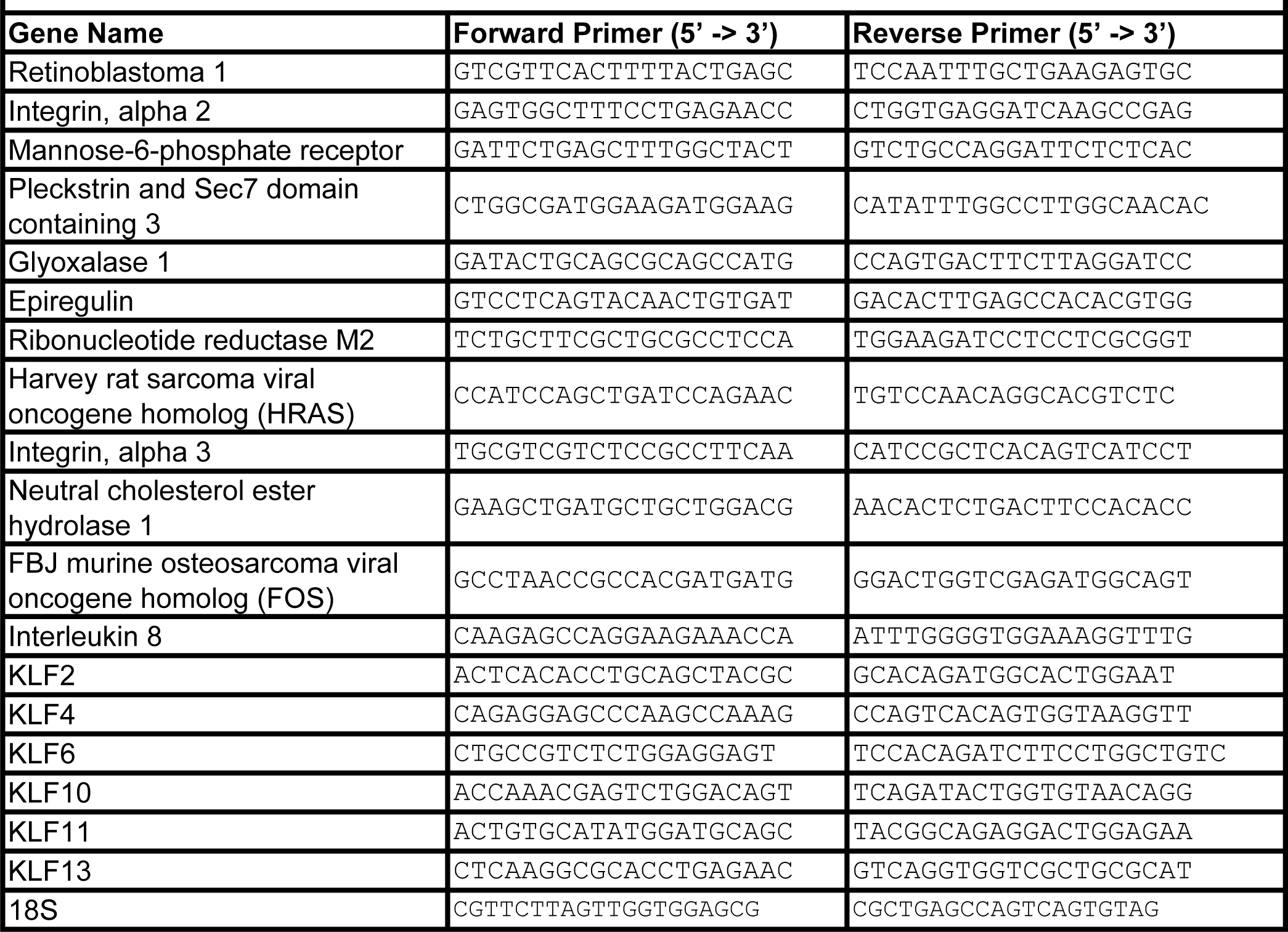
Primers Used for qRT-PCR

### MicroRNA biotinylation

Biotinylation of Cel-miR-67 and miR-106b was performed as described [27]. Mature microRNA for *C. elegans* miR-67 (control) and *H. sapiens* miR-106b (both leading and passenger strand) were from Integrated DNA Tech, Coralville, IA. Leading strand of both Cel-miR-67 and miR-106b was biotinylated at the 3’ end using T4 RNA ligase (pierce RNA 3’ End Biotinylation kit, Thermo Scientific, #20160). Biotinylated RNA was quantified by dot blot on hybond N+. An equal amount of biotinylated mature microRNA (leading strand) was mixed with its respective passenger strand RNA.

### Pull down of biotinylated RNA

Mz-ChA-1 cells were transfected with 50 nM of biotinylated microRNA using Lipofectamine RNAiMAX in triplicate. After 24 hours, cells were lysed (20 mM Tris, pH 7.5, 100 mM KCl, 5 mM MgCl2, 0.3% NP40, 50 U of RNase OUT and complete protease inhibitor) and incubated on ice for 10 minutes. 90% of cell lysate was incubated with streptavidin magnetic beads (New England Biolabs) for 6 hours at 4°C and 10% of cell lysate was used for input RNA. Streptavidin-bead bound RNAs were washed five times with 20 mM Tris, pH 7.5, 0.5 M NaCl and 1 mM EDTA. After pull down, RNA was isolated using *mir*Vana kit. Relative expression of IL8, KLF2 and α Tubulin1A was analyzed by qRT-PCR. 18S was used as a control RNA.

### Immunoblotting

Lysates were probed for KLF2 (Aviva Systems Biology), KLF4 (Cell Signaling), KLF6 (Santa Cruz Biotechnology), KLF10 (abcam), DR5 (Cell Signaling) or actin (Sigma).

### Apoptosis and proliferation

Treated cells were assayed for apoptosis by nuclear morphology, and for cell proliferation by 3 (4,5 dimethylthiazol-2-yl)-2,5-diphenyltetrazolium bromide (MTT) assay, as described [28]. All experiments were repeated at least three times.

## Results

### RNA-Seq to determine miR-106b targets

To determine miR-106b targets, we experimentally manipulated miR-106b levels in human Mz-ChA-1 cholangiocarcinoma cells (**Figure 1A**). Total RNA from each condition, miR-106b and LNA-106b, was sequenced. Messages with decreased counts in miR-106b-transfected samples compared to LNA-106b-transfected samples were defined as repressed by miR-106b, while mRNAs that had increased counts were defined to be increased by miR-106b. More genes were repressed by miR-106b than increased, with 112 mRNAs repressed and 17 increased (**Figure 1B**). Based on the RefSeq sequences, we searched the two data sets—genes with decreased expression and those with increased expression—for the miR-106b seed-binding site. Specifically, we sought perfect 8-mer binding sites and either 7-mer with a match at position 8 (m8) or 7-mer with an A opposite position 1 (A1). The seed sequence of miR-106b is 5’UAAAGUGC and the corresponding 8-mer binding site is 5’GCACUUUA. Of the 112 mRNAs with decreased expression, 81 had either an 8-mer or a 7-mer binding site (72.3%), and 31 had at least one perfect 8-mer binding site (27.7%; **Figure 1C**). Conversely, of the 17 mRNAs with increased expression in cells transfected with miR-106b, only one mRNA contained a 7-mer binding sequence (5.9%) and none had a perfect 8-mer site (**Figure 1C**). The enrichment of seed-binding sites suggested that our technique was effective in detecting miR-106b targets.

**Figure 1.**
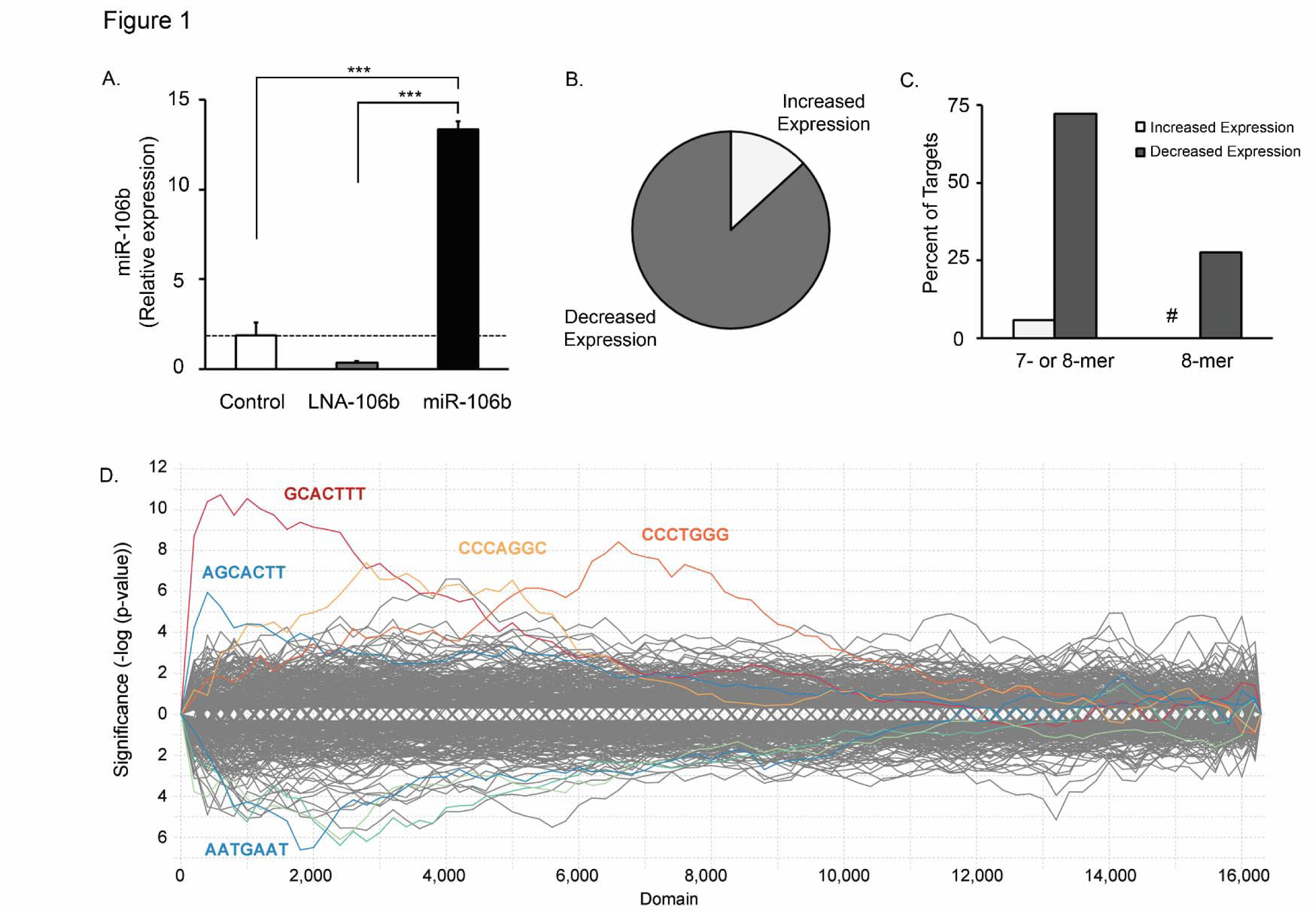
miR-106b targets in cholangiocarcinoma cells predominantly contain a seed-binding site. (A) miR-106b RNA levels were measured by qRT-PCR after transfection of Mz-ChA-1 cells with control LNA (Control), antagonist to miR-106b (LNA-106b), or miR-106b. Expression was normalized to Z30 and plotted as relative level. The dashed horizontal line indicates the mean of miR-106b in Control cells. Data are mean ± SEM for three samples each. *** P < 0.001 using ANOVA with *post hoc* correction. (B) Following RNA-Seq of the LNA-106b and miR-106b samples from panel A, significantly altered transcripts were categorized as increased (13.2%) or decreased (86.8%) by miR-106b. (C) A majority of decreased transcripts contained one or more miR-106b seed-binding sites (7-mer or 8-mer) while only one increased transcript contained a 7-mer binding site. None of the increased transcripts contained an 8-mer binding site (#). (D) All transcripts identified by RNA-Seq were sorted by statistical significance of expression difference between miR-106b and LNA-106b. This sorted list (plotted on the horizontal axis) was then analyzed by SylArray to identify microRNA binding sites that are over-represented, shown by colored traces. Over-representation of seed-binding sites is indicated on the vertical axis above zero, plotted on a log scale. Under-represented sequences are plotted below zero.

### SylArray analysis

The entire gene set was analyzed by SylArray [24] to detect enrichment of microRNA binding sites. We confirmed the enrichment of the miR-106b 7-mer m8 seed-binding site, 5’GCACTTT (red line) in the sequences that were significantly altered by miR-106b (to the left on the plot; **Figure 1D**). Notably, this analysis also revealed enrichment of a 7 nucleotide sequence complementary to nucleotides 3-9 of miR-106b (5’AGCACTT, blue line).

### Sequence determinants of miR-106b targeting

Using previously published data [19], we determined that 76.2% of miR-106b target interactions employed 7 or more consecutive miR-106b bases in the microRNA-mRNA binding hybrid. Because a high proportion of interactions contained at least 7 consecutive bases, we determined which of the miR-106b 7-or 8-mer binding sites were favored in our target gene set. We searched the 112 significant mRNA sequences for all 65,536 possible 8-mer sequences and plotted the frequency of occurrence of each 8-mer (count) versus the number of 8-mers in each bin (**Figure 2A**). Approximately 50% of all possible 8-mers were observed with a frequency of between 1 and 10 occurrences, while 5,020 8-mers were not observed at all (count = 0). Notably, the perfect 8-mer binding site 5’GCACTTTA was present 63 times and the overlapping 8-mer (2-9) was present 81 times. The two most over-represented 8-mers were 5’AAAAAAAA (1,987 times) and 5’TTTTTTTT (830 times). The analogous k-mer data set was developed for all possible 7-mer sequences (16,384 possible 7-mers) within the significant gene set (**Figure 2B**). Again the most common sequences were 5’AAAAAAA and 5’TTTTTTT. Forty-nine 7-mers were absent in the data set. The sequence complementary to bases 1-7 of miR-106b was observed 121 times, the 2-8 sequence was observed 170 times, and the 3-9 sequence was observed 147 times.

**Figure 2.**
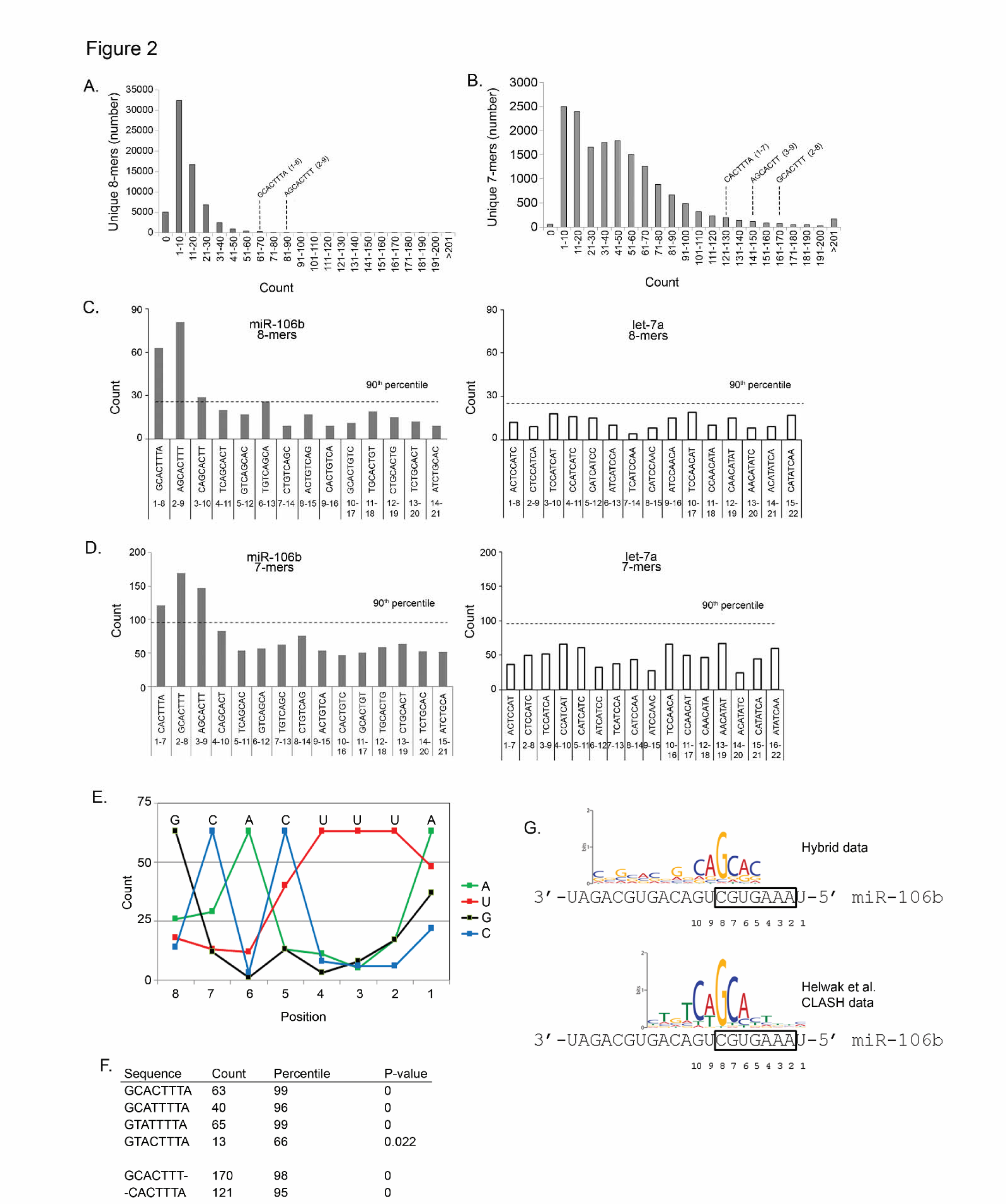
Preferential miR-106b target binding via the 5’ end of the microRNA. (A) Histogram of the distibution of 8-mers found in miR-106b-regulated sequences. The horizontal axis depicts the frequency each 8-mer was observed and the vertical axis represents the number of 8-mers at each count. The bins containing the miR-106b 1-8 and 2-9 perfect binding sequences are indicated. (B) Data are as in panel A except that 7-mers were analyzed. Bins containing the miR-106b 1-7, 2-8, and 3-9 perfect binding sequences are indicated. (C) The count of each possible 8-mer binding site in the miR-106b-regulated sequences is indicated. Sequences are ordered as they occur along miR-106b. The horizontal dashed line represents the count corresponding to the 90th percentile of all 8-mers (i. e., only the top 10 percent occur more frequently). (D) The same plot as in panel C except using 8-mers derived from let-7a. (E) Counts in the miR-106b-regulated sequences of each possible 7-mer along miR-106b are indicated. The horizontal dashed line represents the count corresponding to the 90th percentile of all 7-mers. (F) The same plot as in panel E except using 7-mers derived from let-7a. (G) Single-base substitutions within the 8-base miR-106b binding site were queried for their frequency, compared to the perfect 8-mer (observed 63 times) and plotted as the nucleotide frequency (count) at each position when the other 7 positions were a perfect miR-106b match. For example, the single substitution of a ‘U’ opposite position 5 (forming a G:U wobble) was observed 40 times while the favored ‘C’ (forming a G:C pair) was found 63 times. (H) k-mer analysis of the count of miR-106b binding sites in the miR-106b-regulated gene set compared to 1,000 randomly chosen similarly-sized gene sets. (I) The binding motif for miR-106b is shown over the sequence of the microRNA (antiparallel). Taller letters indicate greater representation of that nucleotide in determining the motif. The upper motif was generated using RNA-Seq data from the current study. The analysis was performed separately on data from Helwak et al., 2010, shown in the lower motif.

Each of the possible miR-106b-complementary 8-mers were tiled from 5’ to 3’ (i.e., 1-8, 2-9, 3-10, etc.). The 8-mer opposite bases 2-9 was somewhat more frequent that the 1-8 complement (**Figure 2C**). A similar plot of the frequency of all complementary 7-mers tiled from 5’ to 3’ showed that over-represented sequences again favored the 5’ end of miR-106b, especially the classical seed (**Figure 2D**). We used an unrelated, control microRNA let-7a; the same plot of tiled 8-and 7-mer sequences showed that none of these sequences were found in the top 10 percentile (**Figure 2C&D**).

To determine the tolerance for single base differences within the seed sequence, we started from a perfect 1-8 sequence and systematically searched for one-off variants. For example, the 1-8 binding site 5’GCACTTTA was observed 63 times while the sequence 5’<u>A</u>CACTTTA occurred 26 times. The sequence 5’GCA<u>T</u>TTTA contains a single base difference opposite position 5 of miR-106b and was observed 40 times (96th percentile). Compare this result to the sequence 5’GCA<u>G</u>TTTA which was observed only 13 times (66th percentile). We plotted the counts for each of these one-off sequences as a function of the position in the miR-106b binding site (**Figure 2E**). The most tolerance was observed opposite position 1 of miR-106b. The next most tolerated substitution was a T at position 5, which would result in a G:U wobble base instead of a G:C pair.

The raw count for each sequence may not indicate over-representation but rather may indicate frequently encountered 8-mers (e.g., 5’AAAAAAAA within the poly-A tail). Thus, we sought to determine the statistical significance of a number of sequences commonly observed in our set of 129 significant genes. This set contained 191 sequences due to some genes having multiple forms. To correct for natural frequency variation, we generated 1,000 additional sets of mRNAs each containing 191 random transcript sequences and compared the frequency of the miR-106b binding sequence and related sequences. The 8-mer sequence 5’GCACTTTA was over-represented in our gene set and this finding was highly statistically significant (p = 0). Similarly, each of the 7-mer sequences was statistically different in our set (p = 0). The two C’s within the miR-106b binding sequence were reassigned as T’s to search for seed-binding sites including a G:U wobble, C5 and C7. The wobble at the 5th position, 5’GCA<u>T</u>TTTA, was highly significant, as was the double wobble, 5’G<u>T</u>A<u>T</u>TTTA. Tolerance for a G:U wobble at position 7 only, 5’G<u>T</u>ACTTTA, was not as highly significant (p = 0.022, **Figure 2F**).

### RNA hybrid analysis

Although the perfect complement is likely preferred, our data indicated that related sequences were also commonly overrepresented. We sought to identify the characteristics of the most thermodynamically stable miR-106b binding site in our significantly-repressed genes. We used RNAhybrid to determine the microRNA:mRNA pairing with the lowest free energy for each repressed target. Next, the predicted miR-106b binding sites were analyzed using MEME Suite to identify the most common motif. The resulting sequence motif includes 6 nucleotides that align with the predicted miR-106b binding site from bases 5-10 (**Figure 2G; upper**). This sequence does not include the triplet U at positions 2-4, possibly because A:T pairs have less favorable free energy than G:C pairs and the query set was limited by the lowest free energy binding site from RNAhybrid.

### Comparison to CLASH data

Supplemental data from [19] included 143 target mRNAs bound by miR-106b, including the sequence at the region of interaction. To validate the miR-106b binding characteristics we have described, we searched for sequence motifs in the CLASH data set using MEME Suite. Within the 143 sequences, 138 contained a sequence highly related to 5’CTGTCAGCACTTTC. This is the complement of the 5’ end of miR-106b from bases 2-14 (with ‘C’ opposite the first position of miR-106b slightly favored over the expected ‘A’). The most conserved region of this meme is 5’CAGCA, complementary to miR-106b bases 6-10 (**Figure 2G; lower**). Comparing the two motifs, the sequence from bases 6-10 is most important with some contribution from flanking bases and tolerance for substitution opposite position 5 (allowing a G:U wobble). The two data sets are mostly in agreement and indicate that binding of miR-106b is anchored by G:C pairing in and near the seed.

### Genome-wide target identification

Next, the identified transcripts in our RNA-Seq data set were plotted by change in expression versus statistical significance (volcano plot). The relative change in expression between miR-106b and LNA-106b samples was plotted as log2 of the fold-change on the horizontal axis against the statistical significance plotted as −log of the p value on the vertical axis (**Figure 3A**). Targets were considered significant if the −log p value was above 3.35 (e.g., p < 4.2 × 10^-4^) and these points are plotted in red for decreased or blue for increased expression. Target gene expression changes of significant genes ranged between −1.16 to −2.22 fold reduced expression and +1.15 to +1.47 fold increased expression. Selected targets are indicated (for a complete list of the significant targets, see **Table 2**). Known target genes RB1 and IL-8 were significantly repressed [10, 29]. Among the novel targets identified in our genome-wide analysis were members of the Kruppel-like factor family, KLF2 and KLF6. These targets were significantly repressed by miR-106b with a p value of 1.72*10^-7^ and 2.85*10^-4^, respectively.

**Figure 3.**
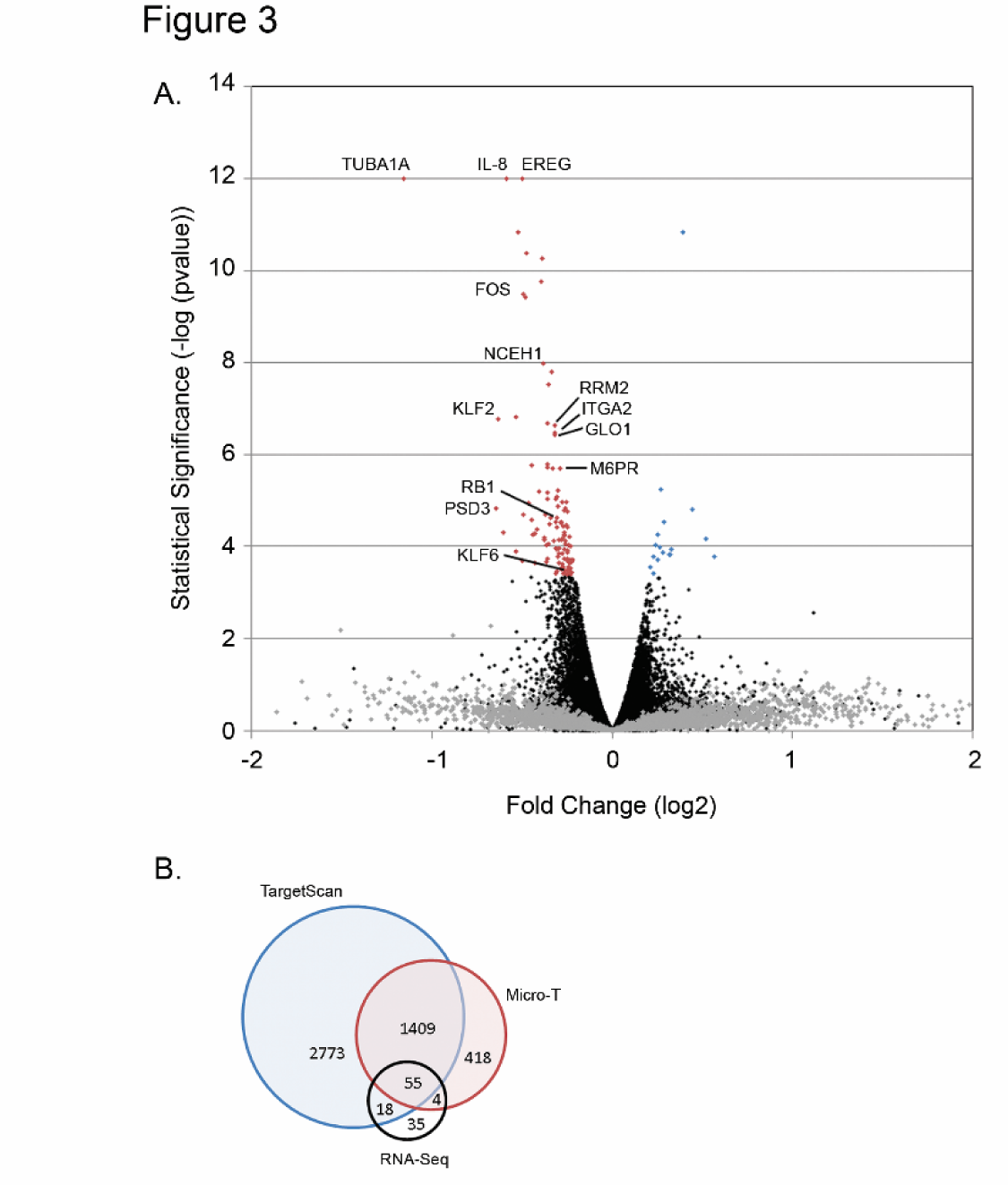
miR-106b target discovery by RNA-Seq. (A) Volcano plot of gene expression by RNA-Seq. miR-106b RNA levels were altered by transfection of Mz-ChA-1 cells with LNA-106b or miR-106b and the resulting differential expression of all genes was evaluated. Transcripts are plotted as log2 of expression fold change on the horizontal axis versus −log of the p value on the vertical axis. Gene expression change was considered significant at −log p > 3.35 and genes with values above this cutoff are indicated by colored points. Significantly altered transcripts were either decreased (red) or increased (blue). Labeled genes are those that have been evaluated for modulation by miR-106b through additional experiments. (B) Venn diagram demonstrating overlap of our RNA-Seq miR-106b target repression results with microRNA target prediction databases TargetScan and Micro-T. Our data contained 35 novel targets not predicted by either program.

**Table 2:**
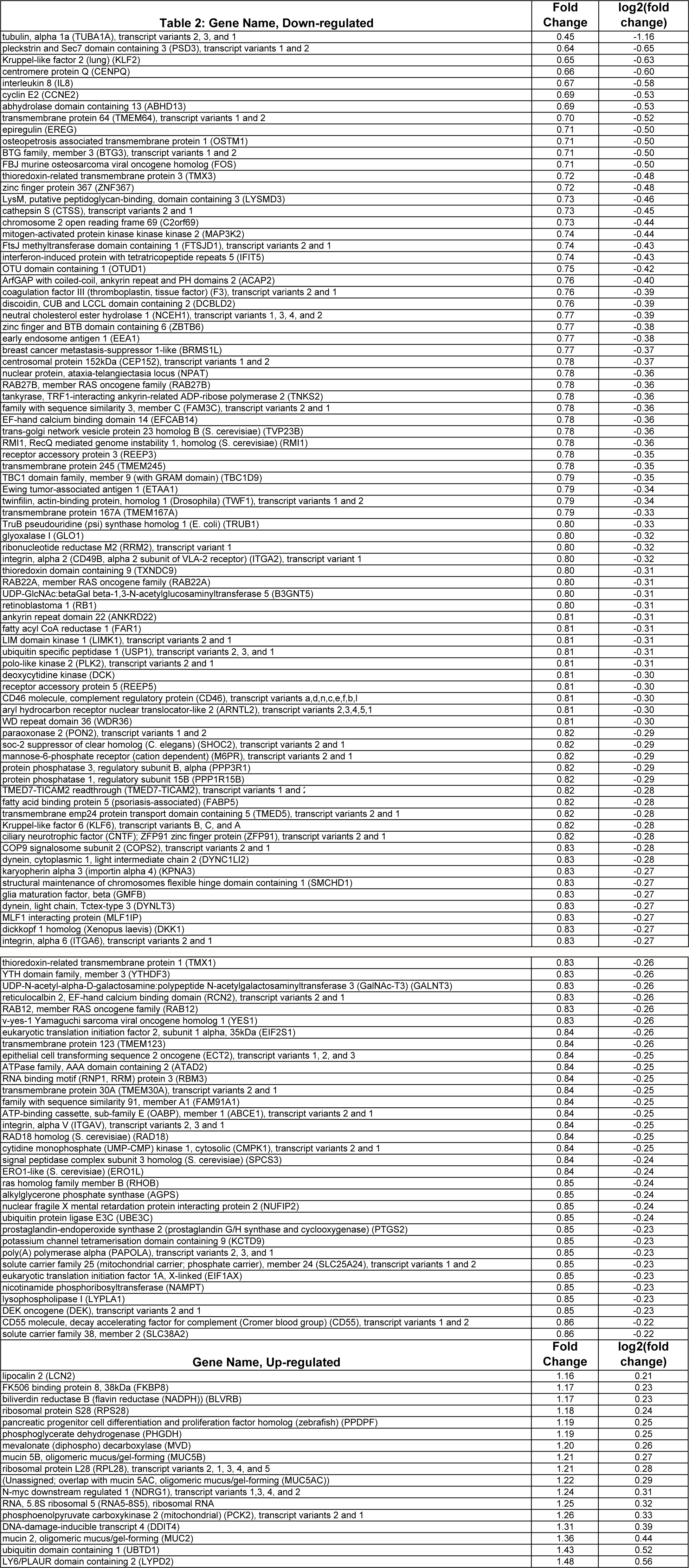
Gene Name, Down-regulated

We compared our experimentally determined set of 112 mRNAs repressed by miR-106b to the genes predicted by TargetScan [30] or Micro-T [31], and compared the two prediction programs to each other as well. Figure 3B shows a Venn diagram of the number of shared targets in these data sets. The majority of genes contained in our data set were predicted by one or both programs, though 35 experimentally identified targets were not predicted by either program (**Figure 3B**). We have also compared our target list with the genes confirmed by CLASH and find that only ERO1L, FAM91A1, and YES1 were identified as miR-106b targets both in our data and that of Tollervey and colleagues [19].

### Target validation

We sought to verify that miR-106b levels altered expression of target genes identified by RNA-Seq. We transfected Mz-ChA-1 cells with negative control LNA, miR-106b, or LNA-106b and isolated total RNA. Transcript levels of selected targets were determined by quantitative RT-PCR and normalized to 18S rRNA. Targets confirmed by RT-PCR include EREG, RRM2, ITGA2, RB1, GLO1, M6PR, and PSD3. We also evaluated non-target negative controls ITGA3 and HRAS and found no change by LNA-106b compared to negative control LNA. (**Figure 4A**). NCEH1 was a target identified by RNA-Seq which had a trend towards increased mRNA expression upon miR-106b antagonism, but was not statistically significant (p = 0.07). We did not observe a change in mRNA level in RNA-Seq target FOS by RT-PCR. Not all miR-106b targets will be identified by the current method, specifically those that do not have a decrease in mRNA levels. We have not exhaustively tried to determine the identity of such targets, but did recognize the TRAIL death receptor DR5 as a predicted miR-106b target by TargetScan. Because miR-106b is clustered with miR-25 and miR-25 regulates cell death by targeting (DR4) [4], we tested whether miR-106b decreased the functionally-related DR5 protein. Transfection with miR-106b decreased DR5 protein levels (**Figure 4B**), potentially acting to complement miR-25-mediated DR4 repression and promote TRAIL resistance.

**Figure 4.**
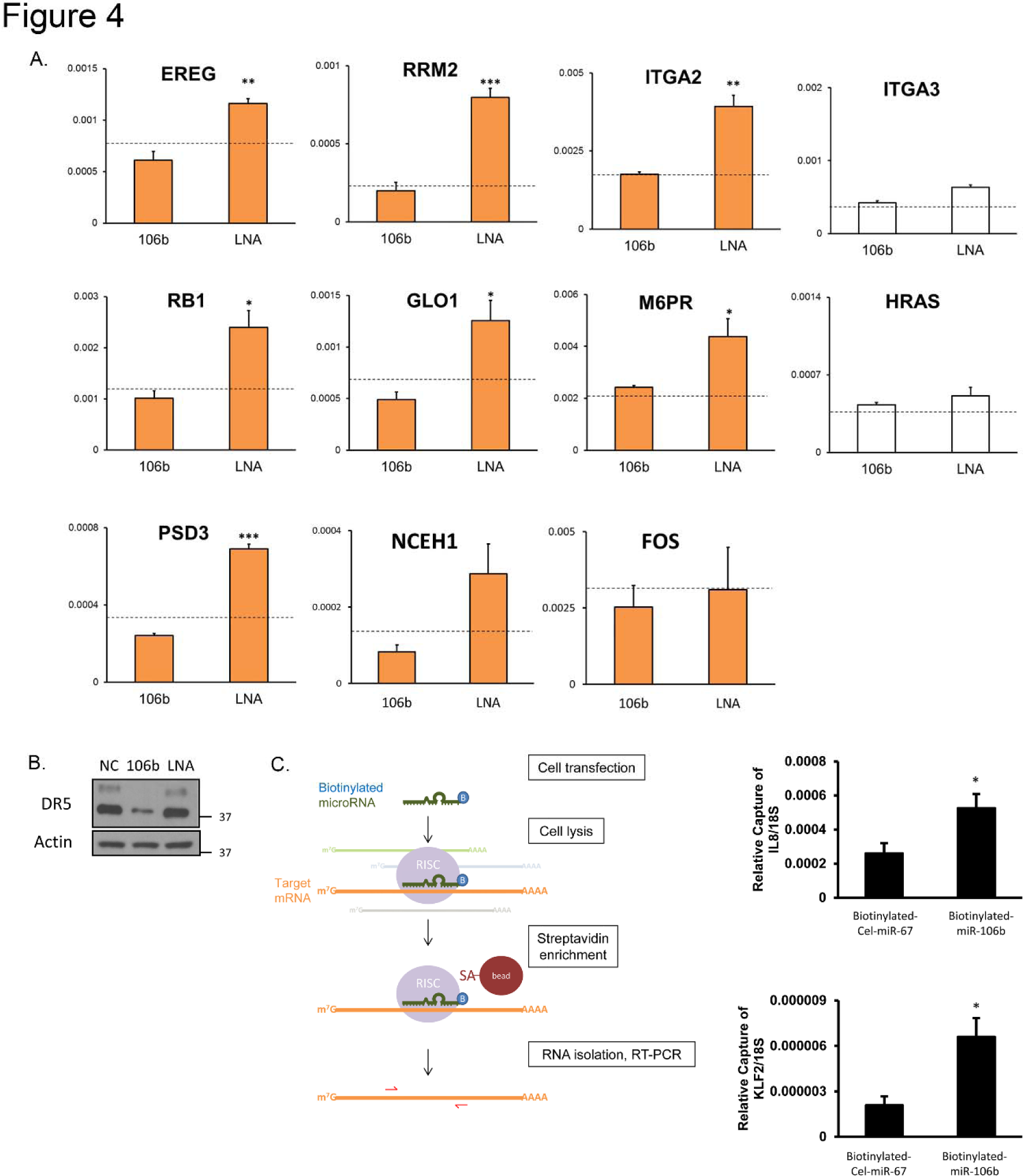
RNA-Seq target validation. (A) qRT-PCR for nine candidate targets from RNA-Seq. Relative expression is significantly increased for LNA-106b compared to miR-106b in six of the genes. There was a trend towards increased expression for LNA-106b compared to miR-106b for NCEH1 but it was not statistically significant (p = 0.07). FOS showed no significant change in expression by qRT-PCR. Dotted line represents expression level for scrambled control LNA. ITGA3 and HRAS are non-target negative control genes which show no significant expression change. (B) Immunoblot showing transfection with miR-106b decreased the functionally-related DR5 protein levels. (C) Schematic of experimental design for capture of mRNA targets using biotinylated microRNA. Briefly, Mz-ChA-1 cells were transfected for 24 hours with either mature human miR-106b or *C. elegans* miR-67 which had been biotinylated. Cells were lysed and incubated with streptavidin-bound beads to capture biotinylated microRNA and associated mRNAs. Total RNA was isolated and relative expression of target mRNAs KLF2 and IL-8 was measured for enrichment by qRT-PCR. 18S was used as a control RNA. * p < 0.05; ** p < 0.01; *** p < 0.001; using ANOVA with *post hoc* correction.

We used biotinylated miR-106b in order to validate targets by affinity binding and capture. Mz-ChA-1 cells were transfected with either biotinylated miR-106b or biotinylated *C. elegans* miR-67 (Cel-miR-67) as a control. Biotin-bound RNA was isolated and RT-PCR was performed for IL-8 and KLF2. We observed significant enrichment of IL-8 and KLF2 transcripts with biotinylated miR-106b pulldown versus control Cel-miR-67 pulldown (**Figure 4C**).

### miR-106b targets multiple KLF family members

Based on the observed decrease in KLF2 and KLF6 in the RNA-Seq data set, we tested the effect of miR-106b on additional KLF family members. KLFs represent a large family of transcription factors of which many act as tumor suppressors and are often down-regulated in cancer [32, 33]. We confirmed that KLF2 and KLF6 mRNAs were regulated by miR-106b and found other KLF family members KLF4, KLF10, KLF11 and KLF13 to have increased expression when cells were transfected with LNA-106b (**Figure 5A**). Additionally, we examined the effects of miR-106b on protein expression of KLFs by immunoblot in Mz-ChA-1 cells. We observed decreased expression of KLF2, KLF4, KLF6, and KLF10 after 24 hours of transfection with miR-106b compared to negative control LNA and a slight increase in expression with LNA-106b (**Figure 5B**).

**Figure 5.**
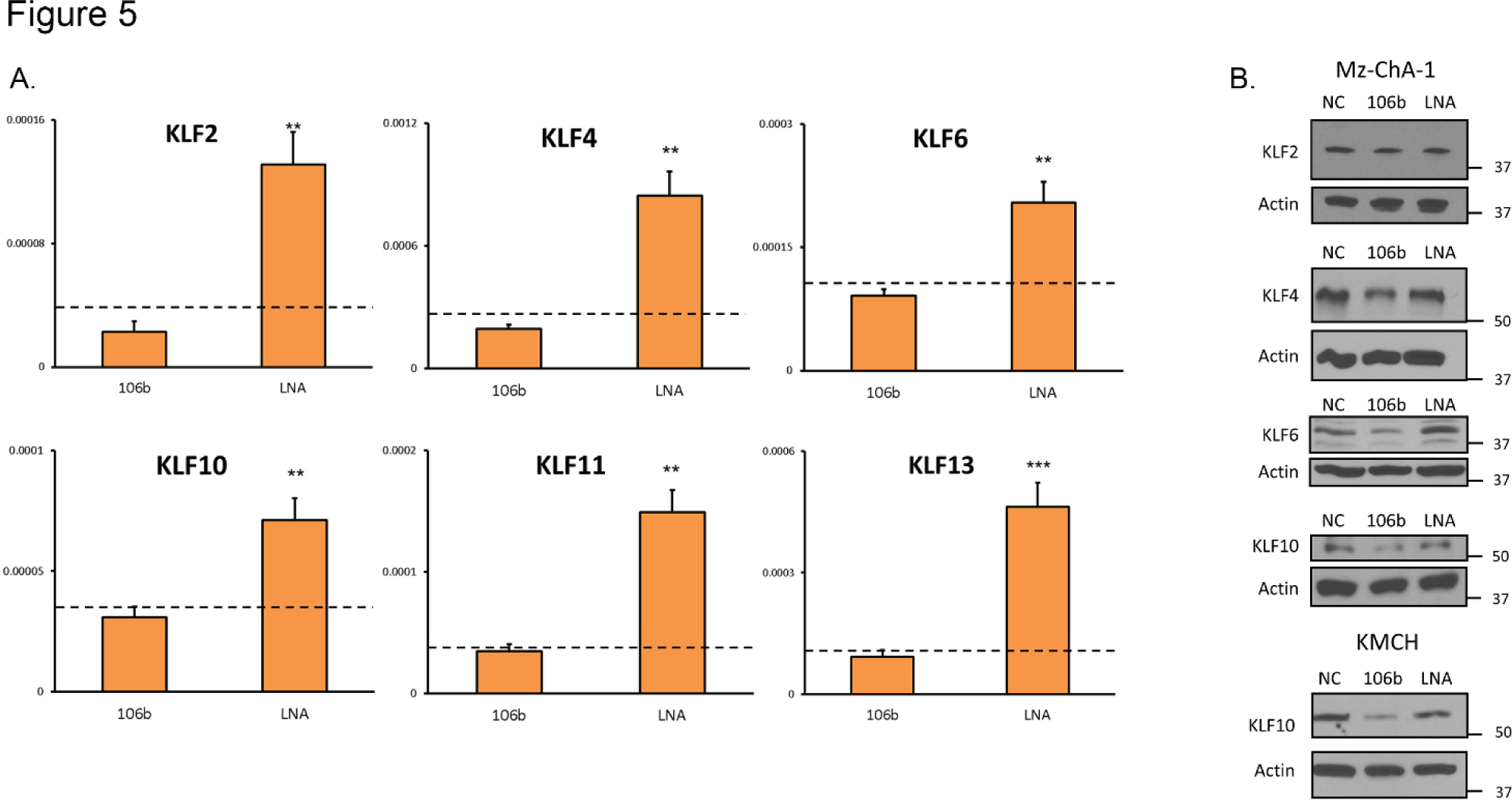
miR-106b regulates multiple KLF family members. (A) qRT-PCR for RNA-Seq candidate miR-106b targets KLF2 and 6 confirmed regulation at the RNA level. mRNA expression was increased by miR-106b antagonism with LNA-106b compared to miR-106b. Additional KLFs 4, 10, 11, and 13 were evaluated and showed the same pattern of expression increase with LNA-106b antagonism compared to miR-106b treatment. (B) Immunoblots showing regulation of KLFs 2, 4, 6 and 10 by miR-106b at the protein level. 24 hour transfection of Mz-ChA-1 or KMCH cells with control LNA, miR-106b, or LNA-106b led to decrease in KLF protein expression by miR-106b compared to control LNA or LNA-106b. Actin was probed as a loading control. * p < 0.05; ** p < 0.01; *** p < 0.001; using ANOVA with *post hoc* correction.

### Proliferation

To investigate the role of miR-106b in proliferation of cholangiocarcinoma cells, we assessed the change in cell number over time using the MTT assay. Mz-ChA-1, KMCH, and BDEneu cholangiocarcinoma cells were transfected with miR-106b, LNA-106b or negative control LNA for 24 hours. Cells were then allowed to grow for up to 72 hours. We did not observe any significant difference in cell proliferation upon alteration of miR-106b levels (**Figure 6**). To eliminate the possibility of a long-term effect on cell growth we repeated the assay over a one week course in Mz-ChA-1 cells and again observed no change in proliferation (data not shown).

**Figure 6.**
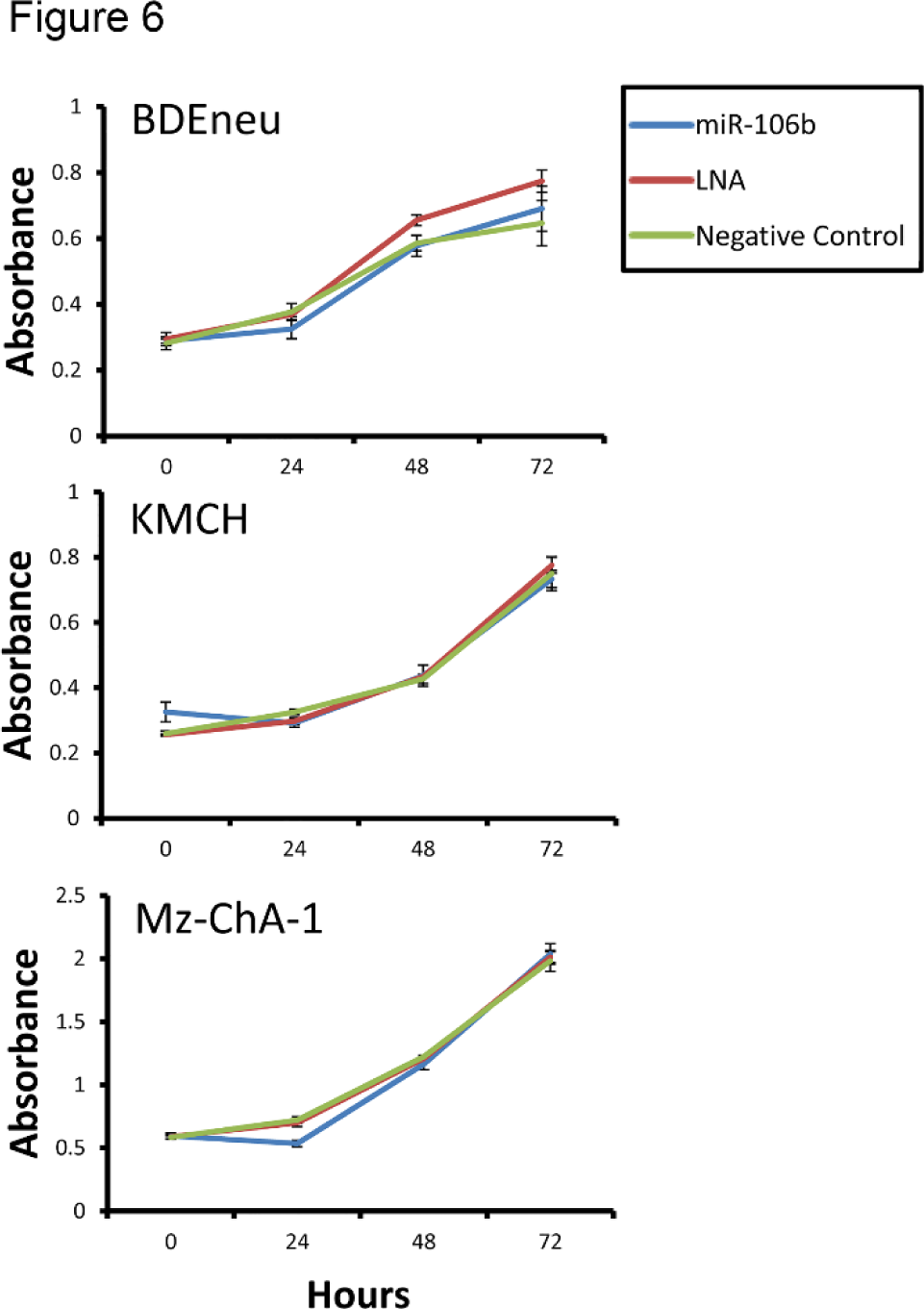
miR-106b does not affect proliferation in cholangiocarcinoma cells. We tested the effect of miR-106b on proliferation in BDEneu, KMCH, and Mz-ChA-1 cells by MTT assay. After 24 hour transfection with control LNA, miR-106b or LNA-106b, cells were allowed to grow for up to 72 hours and cell number was measured by absorbance read at 540 nm. We observed no significant difference in cell growth for any cell line. Signal represents the mean (n?=?4) +/− SEM.

### miR-106b protects against apoptosis

KLF2, KLF6 and KLF10 were all regulated by miR-106b and are known to promote apoptosis [34,35,36]. Our data demonstrated additionally that DR5, a pro-apoptotic death receptor, was regulated by miR-106b. Thus, we reasoned that miR-106b may protect against apoptosis in cholangiocarcinoma cells. H69, KMCH, and Mz-ChA-1 cells were transfected with miR-106b, LNA-106b or negative control followed by treatment with either TRAIL or staurosporine to induce cell death. We observed a decrease in apoptotic nuclei by DAPI staining upon transfection with miR-106b and an increase in apoptotic nuclei upon transfection with LNA-106b (**Figure 7**). Thus, miR-106b acts in part to protect cholangiocarcinoma cells from apoptosis.

**Figure 7.**
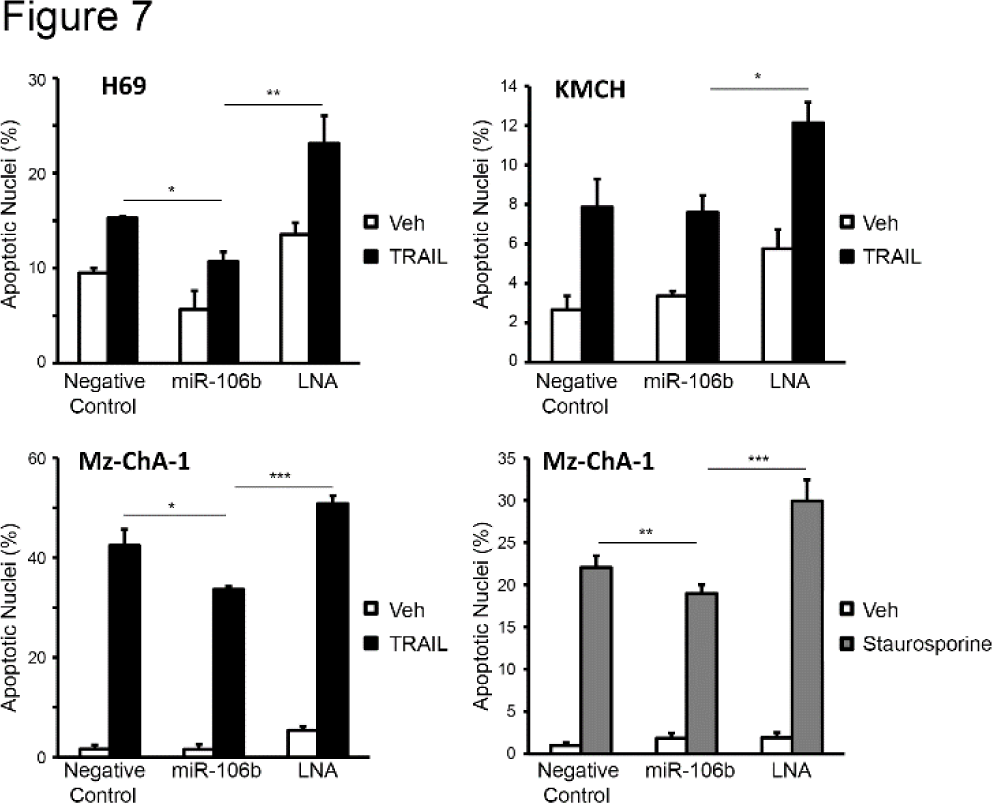
miR-106b protects against TRAIL-or staurosporine-induced apoptosis in cholangiocarcinoma cells. H69, KMCH or Mz-ChA-1 cells were transfected with control LNA, miR-106b or LNA-106b for 24 hours followed by treatment with either TRAIL or staurosporine to induce apoptosis. We observed a decrease in apoptotic nuclei by DAPI staining upon transfection with miR-106b and an increase in apoptotic nuclei upon transfection with LNA-106b. DAPI-positive nuclei were counted and expressed as a percent of total nuclei. Data are a mean of three experiments +/− SEM. * p < 0.05; ** p < 0.01; *** p < 0.001; using ANOVA with *post hoc* correction.

## Discussion

The data presented in this manuscript relate to mRNAs regulated by miR-106b in cholangiocarcinoma cells. A number of cancer types over express miR106b, making a target gene set of potential value to tumors beyond cholangiocarcinoma. The principle findings reported here show: (i) miR-106b repressed 112 mRNA targets; (ii) most target genes contained a 7‐ or 8-mer seed-binding site; (iii) multiple KLF family proteins are targeted by miR-106b; and (iv) miR-106b promoted tumor cell survival in cholangiocarcinoma cells. Each of these findings will be discussed below.

Our study has revealed 112 mRNAs that were negatively regulated by miR-106b. Some known miR-106b-regulated genes (e.g., RB1, IL-8, F3, YES1, FAM91A1, and ERO1L) were identified and several novel targets were uncovered as well. Our experiments did not further investigate a functional role for these known targets. Not all previously-identified miR-106b-regulated genes were significantly altered in our study. Specifically, we did not observe any change in the mRNA levels of PTEN, E2F1, or BCL2L11 (Bim). A lack of change in expression of these mRNAs may reflect cell-line specific differences in microRNA targeting, changes in mRNA levels below the threshold of detection, or post-transcriptional effects that do not change mRNA levels. Previously unknown genes that were regulated at the mRNA level include KLF family members, which are discussed below.

MicroRNA binding depends on sequence complementarity. The degree of complementarity and length of consecutive bases can vary, resulting in refinement of rules of binding [37] and definition of new types or classes of microRNA:mRNA interactions [19]. Classes I-III of interactions involve complementarity within the seed region. Class IV interactions show complementarity in a more central region, described as centered pairing [18]. Finally, Class V interactions exhibited distributed pairing, where discrete continuous regions of complementarity were not observed [19]. These data are consistent with a role for microRNAs in regulating expression of mRNAs based on sequence complementarity but not strictly limited to seed pairing. Perhaps not surprisingly, the GC content of microRNA binding motifs, representing the average or commonly identified binding site over many mRNAs, was higher than the GC content of microRNA seed regions in general [19]. Thus, binding energy may be more important than binding position, a concept incorporated into the microRNA target prediction algorithm RNAhybrid [25].

Over 70% of mRNAs that were decreased contained either a 7-mer or 8-mer miR-106b binding site. Analysis of all 8-mers or 7-mers in miR-106b-regulated sequences demonstrated that the sequences at or near the seed, including up to nucleotide 10, were over-represented. This over-representation of the 7-mer and 8-mer sequences was highly statistically significant when compared to the expected distribution of the same sequences in a thousand random gene sets of the same size. The seed sequence tolerated a ‘U’ in the place of ‘C’ (resulting in G:U wobble pairing) opposite positions 5, 7, or both. Two hydrogen bonds connect the G:U pair while three hydrogen bonds stabilize the G:C pair, suggesting there may be a thermodynamic cost to miR-106b binding sites with G:U wobbles. Alternatively, the exocyclic amino group of ‘G’ is available for additional interactions when ‘G’ is paired with ‘U’ [38] allowing for the possibility of compensatory stabilizing hydrogen bonds to functional groups within the RISC polypeptides.

Based on both the CLASH dataset [19] and our own, we found that the sequence 5’U<u>CAGCAC</u>U represents an ‘average’ sequence motif serving as a binding site for miR-106b, with the best evidence for the central 6-mer (underlined, complementary to miR-106b based 5-10). While this is the average binding sequence, the most prevalent 7-mer was 5’GCACUUU and the most common 8-mer was 5’AGCACUUU. The difference between the average and the most prevalent sequences is that the average (in our data set) was determined using the single-most-stable predicted binding site, as identified by RNAhybrid. Such a filter will bias against the triplet UUU. Still, the agreement between our average binding motif and the motif generated from CLASH data where this potential bias is not relevant suggest that this filter is not unreasonable. Overall, we find good evidence that binding favors complementarity near the 5’ end of the microRNA, consistent with the seed model, as well as evidence that the sequence tolerates a slight shift toward the center of miR-106b.

Comparison of our data set of modulated genes and that obtained by CLASH showed a striking near-absence of overlap in the mRNAs identified. Indeed, of the 143 mRNAs in the CLASH set and 112 genes down regulated in our experiment, only three mRNAs were on both lists: ERO1L, YES1, and FAM91A1. None of these contain an 8-mer binding site. YES1 and FAM91A1 each have a single copy of the 7-mer-m8 binding site and YES1 has an additional 7-mer-A1 binding site. The cell lines used in the two studies are very different, Flp-In T-REx 293 embryonic kidney-derived cells versus Mz-ChA-1 biliary tract cancer-derived cells. The techniques used are also different. Finally, data from the CLASH study reflect steady-state interactions of all detected microRNAs and their targets, while the current study assessed acute changes to mRNA targets after manipulation of only miR-106b. It is likely that different cell types will have a different microRNA target landscape, and that identification of these targets by several methods will allow a detailed understanding of mRNA regulation and binding by microRNAs.

An important finding in the current study was the coordinated regulation of multiple KLF family members. The seventeen members of the KLF family of transcription factors are involved in a diverse range of biological functions and aberrations can lead to disorders such as cardiovascular and respiratory disease, obesity, inflammatory diseases and cancer [39]. Many KLF family members have been implicated in some aspect of cancer cell biology including growth, apoptosis, differentiation, and migration [40]. We have revealed regulation of six KLF members by miR-106b in our study. Two members, KLF2 and KLF6, were significantly repressed genes in our RNA-Seq experiment. Additionally, KLF4, KLF10, and KLF11 were somewhat near the cutoff for significance (p = 0.024, 0.038, and 0.032 respectively), while KLF13 mRNA in the RNA-Seq data did not suggest regulation (p = 0.81). These six KLFs were demonstrated to be miR-106b-responsive genes by qRT-PCR. In particular, the result for KLF13 was surprising as this gene was included as a control under the expectation it would not be responsive to miR-106b. The coordinated modulation of these six family members may indicate a functional aspect of miR-106b biology that was not previously appreciated. Each of the KLFs regulated by miR-106b in our study has tumor suppressive function in one or several cancers. KLF2 has been shown to induce apoptosis and to be a tumor suppressor in prostate and breast cancer cell lines as well as in xenografted mice [33]. In pancreatic cancer cells, KLF2 expression is decreased and its enforced expression leads to inhibition of growth and metastasis [41]. KLF4 regulates proliferation and differentiation of lung cancer cells and its deletion in a mouse is enough to generate tumors [42]. KLF6 reduced tumorigenic features in osteosarcoma cells [43] and its expression is reduced in human and mouse prostate cancer [44]. Loss of KLF6 expression results in increased liver mass, decreased cyclin-dependent kinase inhibitor p21, and correlated with low p21 in liver cancer [45]. Mice deficient in KLF10 exhibit increased skin tumorigenesis when exposed to carcinogens [46]. Knockdown of KLF11 in leiomyoma cells leads to increased proliferation and its expression is lower in tumor tissue compared to normal [47]. KLF13 is shown to repress anti-apoptotic Bcl-2 in acute lymphoblastic leukemia [48]. These varied and overlapping anti-tumorigenic features of KLFs in cancer make them attractive potential targets for future study in cholangiocarcinoma.

Resistance to apoptosis is a characteristic of cholangiocarcinoma cells. Many KLFs are implicated in regulation of apoptosis in tumor cells but their role in cholangiocarcinoma cell death is unknown. We have shown regulation of six KLFs by miR-106b. Overexpression of KLF2 in hepatocellular carcinoma cell lines led to increased cell death [49]. KLF6 has dual roles in apoptosis as its wildtype form is pro-apoptotic [50] while the splice variant SV1 which is overexpressed in cancer [51] is anti-apoptotic. KLF10 promotes cell death in human leukemia cells through upregulation of pro-apoptotic proteins Bim and Bax [36]. In our study, antagonism of miR-106b with LNA led to increased apoptosis sensitivity in cholangiocarcinoma cells. In part, this effect could be through derepression of pro-apoptotic KLFs. Targeting of miR-106b to increase cholangiocarcinoma cell sensitivity to apoptosis is a potential future strategy.

In summation, we report a landscape of miR-106b responsive genes in cholangiocarcinoma cells. Several tumor suppressive members of the KLF family of transcription factors were revealed to be modulated by miR-106b. And finally, miR-106b is protective against apoptosis in cholangiocarcinoma cells.

